# Aromatic cage-directed azide-methyllysine photochemistry for profiling non-histone interacting partners of the MeCP2 methyl-CpG binding domain

**DOI:** 10.1101/2025.11.15.688584

**Authors:** Jyotirmayee Padhan, Babu Sudhamalla

## Abstract

Methyl-CpG binding protein 2 (MeCP2) is a well-characterized DNA methylation reader that has recently been shown to interact with methylated histones. MeCP2 recognizes tri-methylated lysine 27 on histone H3 (H3K27me3) through an aromatic cage within its methyl-CpG binding domain (MBD), and this interaction contributes to the regulation of MeCP2 target genes. However, whether MeCP2 can bind methylated non-histone proteins remains unknown. In this study, we sought to identify novel, aromatic cage-dependent non-histone interacting partners of MeCP2 using an unnatural amino acid-based photocrosslinking chemoproteomic strategy. We engineered the aromatic cage of MeCP2-MBD by incorporating the photocrosslinkable amino acid 4-azido-L-phenylalanine (AzF). The AzF-incorporated MeCP2-MBD variants efficiently crosslinked with histone H3 and with methylated histone marks such as H3K4me3 and H3K27me3. MeCP2-MBD-F142AzF was subsequently used to crosslink proteins from HEK293T cell lysates, and the enriched complexes were analyzed by mass spectrometry. This approach identified numerous previously unrecognized interacting partners of MeCP2, many of which are involved in key cellular processes including chromatin regulation, RNA processing, translation, and metabolism. These findings reveal that MeCP2 functions extend beyond DNA methylation reading and transcriptional repression, highlighting a broader role for MeCP2 in coordinating cellular homeostasis.

## Introduction

Methyl-CpG binding protein 2 (MeCP2) is a reader of DNA methylation and the founding member of methyl CpG binding domain (MBD) protein family (1,2). It was initially discovered in the year 1992 in Adrain Bird’s laboratory while attempting to isolate factors responsible for binding unmethylated DNA (3). MeCP2 is largely an intrinsically disordered protein but has several functional domains like the MBD, transcription repressor domain (TRD), NCoR-interacting domain (NID) and three AT hook motifs (4,5) (Figure 1A). Among these, only the MBD comprises of specific secondary structural characteristics and adopts a wedge-like shape consisting of two α-helices and three anti-parallel β-strands (6). The less structured TRD only becomes ordered once the MeCP2 protein is bound to the DNA (7–9). TRD of MeCP2 recruits corepressor proteins like histone deacetylases (HDACs) and Sin3A complex to mediate chromatin repression (10,11). While, the NID interacts with transducin-beta like 1 (TBL1) and TBL1-related (TBLR1) which are the core component of NCoR1/2 complex (12,13). In addition, the AT-hook motifs facilitates MeCP2’s interactions with chromatin by recognizing the AT-rich DNA sequences (9,14,15).

**Figure 1.**
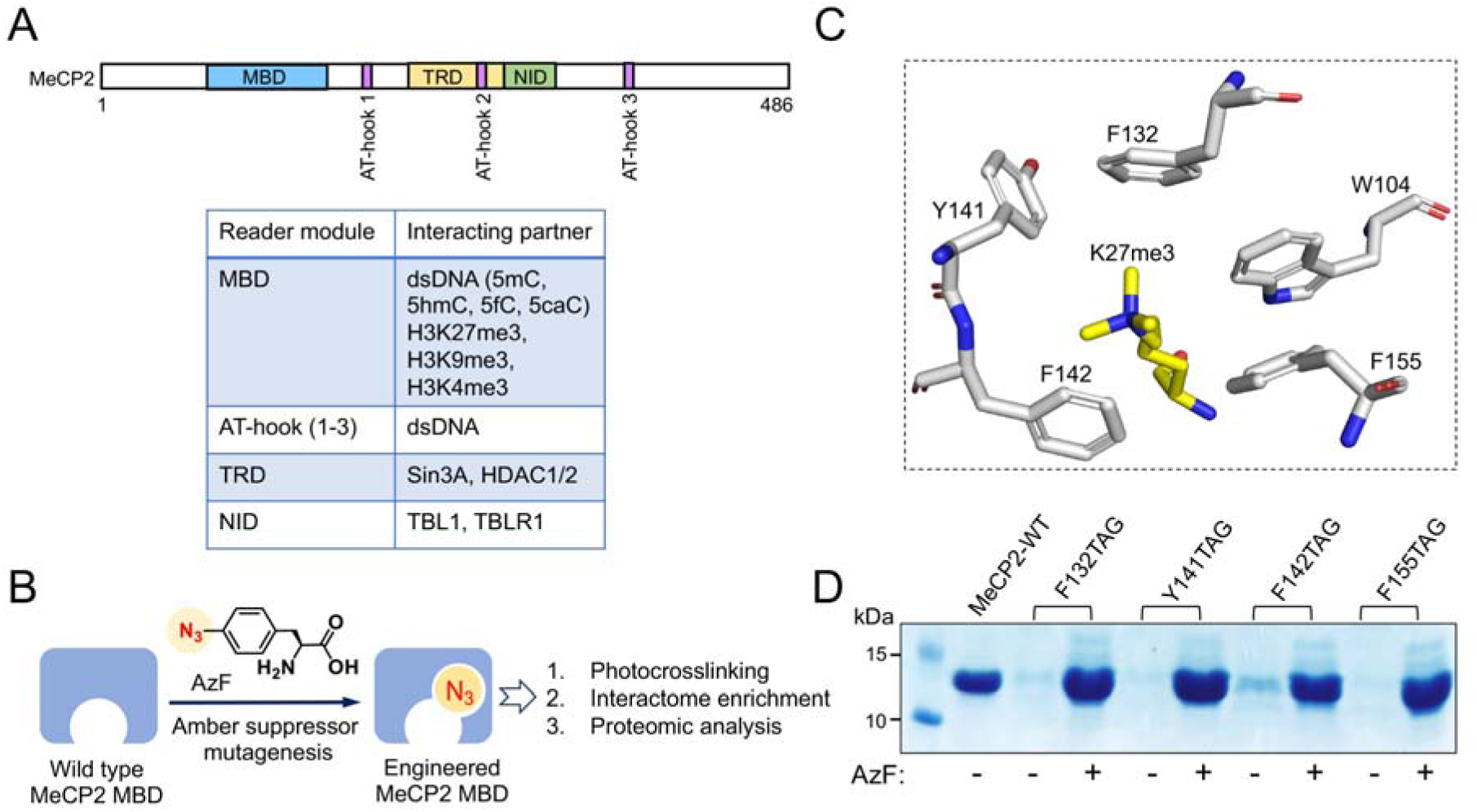
Engineering MeCP2-MBD with 4-azido-L-phenylalanine (AzF). (A) Schematic representation of the domain organization of MeCP2, showing the MBD, TRD, NID and three AT hook motifs along with their known interacting partners. (B) Schematic depicting the engineering of MeCP2-MBD with photocrosslinkable amino acid AzF. (C) Close-up view of docked complex of MeCP2-MBD with H3K27me3 peptide, highlighting the aromatic cage used for residue selection. (D) Coomassie-stained SDS-PAGE gel showing overexpression of MeCP2-MBD-WT and AzF incorporated mutant proteins.

Although ubiquitously expressed, MeCP2 is predominantly a neuronal protein and plays essential role in neurodevelopment (16,17). The expression of MeCP2 increases following neurogenesis and reaches high levels in mature neurons, remaining high throughout adulthood, underscoring the importance of MeCP2 in neural differentiation as well as in adult brain (16,18,19). Studies suggests that MeCP2 regulates expression of neural markers and genes involved in cell fate determination during neurogenesis (19–21). MeCP2 act as a classical transcription regulator and chromatin architect, functioning to reduce aberrant transcriptional events (19,22). In the neuronal genome MeCP2’s concentration is near equivalent to histone octamer and often antagonize histone H1 in binding to promote tertiary chromatin structure formation and compaction (1,19,22–24). MeCP2 also contributes in chromatin looping, chromocenter formation and heterochromatin reorganization during neuronal differentiation (19,22). MeCP2 broadly act as a chromatin organizer, affecting global chromatin landscape and fine-tuning neuronal gene expression (19,22).

Mutation or dysregulation of MeCP2 cause severe neurodevelopmental disorders like Rett syndrome and MeCP2 duplication syndrome (MDS), which manifests as developmental delays, cognitive impairment, learning and intellectual disability, autism spectrum disorder, and seizures (25). In addition to its well-established role in neurodevelopment, MeCP2 has also been implicated in various cancers. Investigation of The Cancer Genome Atlas (TCGA) database revealed that MeCP2 is frequently amplified and overexpressed across numerous human cancer types, including breast, lung, cervical, liver, and stomach (26). MeCP2 could functionally substitute activated RAS and induce the major growth factor signalling pathways, such as MAPK and PI3K in human mammary epithelial cell (HMEC) line (26). Moreover, the overexpression of MeCP2 in breast cancer has direct correlation with poorer patient survival (27). Mechanistically, MeCP2 promotes breast cancer cell proliferation and inhibits apoptosis by facilitating ubiquitination mediated P53 degradation (27). Collectively, emerging evidence identifies MeCP2 as a potential oncogene, acting through diverse molecular mechanism to promote cancer progression and tumorigenesis (28).

Despite extensive studies on MeCP2’s role as a DNA-methylation reader and chromatin organizer, the molecular mechanisms by which it exerts such diverse regulatory functions are not completely understood. MeCP2 interacts with a range of chromatin modifiers, transcriptional regulators, and signaling proteins to modulate gene expression (9,29). However, the full spectrum of its interacting partners is yet to be comprehensively characterized. Our recent work demonstrated that the MBD domain of MeCP2 contains an aromatic cage capable of recognizing histone methylation, thereby influencing gene expression and chromatin association (30). These findings suggest that MeCP2 might also engage in methylation-dependent interactions with non-histone proteins, but identifying such interactions is challenging because they are often transient and weak.

To address this challenge, we turned to strategies developed for studying methyllysine-mediated interactions. Methyllysine sites in proteins are recognized by a variety of reader domains that mediate protein-protein interactions critical for cellular regulation. Previous work by Islam and colleagues demonstrated that chromodomains, canonical methyllysine readers, can be engineered to incorporate the photoactive amino acid 4-azido-L-phenylalanine (AzF) via amber suppressor mutagenesis, enabling selective binding and photocrosslinking of methylated proteins in mammalian cells (31). Building on this strategy, we engineered the methyllysine-binding pocket within the MeCP2 MBD to incorporate the photocrosslinkable unnatural amino acid AzF (Figure 1B), allowing us to capture transient and weak interacting partners of MeCP2. Using this approach, we uncovered numerous previously unrecognized interacting partners, revealing that the functional repertoire of MeCP2 extends far beyond classical transcriptional regulation.

## Results

### Engineering the MeCP2-MBD with photocrosslinking amino acid

In our previous study, we characterized the methyllysine-binding pocket within the MBD of MeCP2 and identified five aromatic residues W104, F132, Y141, F142, and F155 that form an aromatic cage responsible for recognizing the trimethyllysine of H3K27me3 (30). To capture the transient non-histone interactome of MeCP2, we aimed to engineer the MeCP2-MBD with the photocrosslinkable unnatural amino acid 4-azido-L-phenylalanine (AzF) without disrupting binding to its cognate ligand. Guided by the docked complex of MeCP2-MBD with the H3K27me3 peptide (Figure 1C), we targeted tyrosine and phenylalanine residues within the cage because of their structural similarity to AzF. Consequently, four photocrosslinkable MeCP2-MBD variants F132AzF, Y141AzF, F142AzF, and F155AzF were generated via amber codon (TAG) mutagenesis using the evolved orthogonal *Methanococcus jannaschii* TyrRS-tRNA_CUA_^Tyr^ pair (32). The MeCP2-MBD variant with AzF in the amber codon position was synthesized in *E. coli* cells when AzF was included in culture medium. All variants expressed well and were purified to high homogeneity (Figure 1D).

### Photocrosslinking of engineered MeCP2-MBD variants with histone substrates

To assess the photocrosslinking efficiency of the engineered MeCP2-MBD-AzF variants, we performed photocrosslinking experiments using endogenous histones isolated from HEK293T cells as substrates. The engineered MeCP2-MBD-AzF variants were incubated with the histones and then irradiated with 365 nm UV light (Figure 2A). Crosslinked products were enriched using Ni-NTA beads and analyzed by immunoblotting with an anti-H3 antibody. Our results indicate that all MeCP2-MBD-AzF variants successfully crosslinked with histone H3, as evidenced by the appearance of high-molecular-weight bands upon UV irradiation (Figure 2B). Among the variants, Y141AzF displayed noticeably lower crosslinking efficiency with H3. No crosslinked products were detected in the absence of UV exposure.

**Figure 2.**
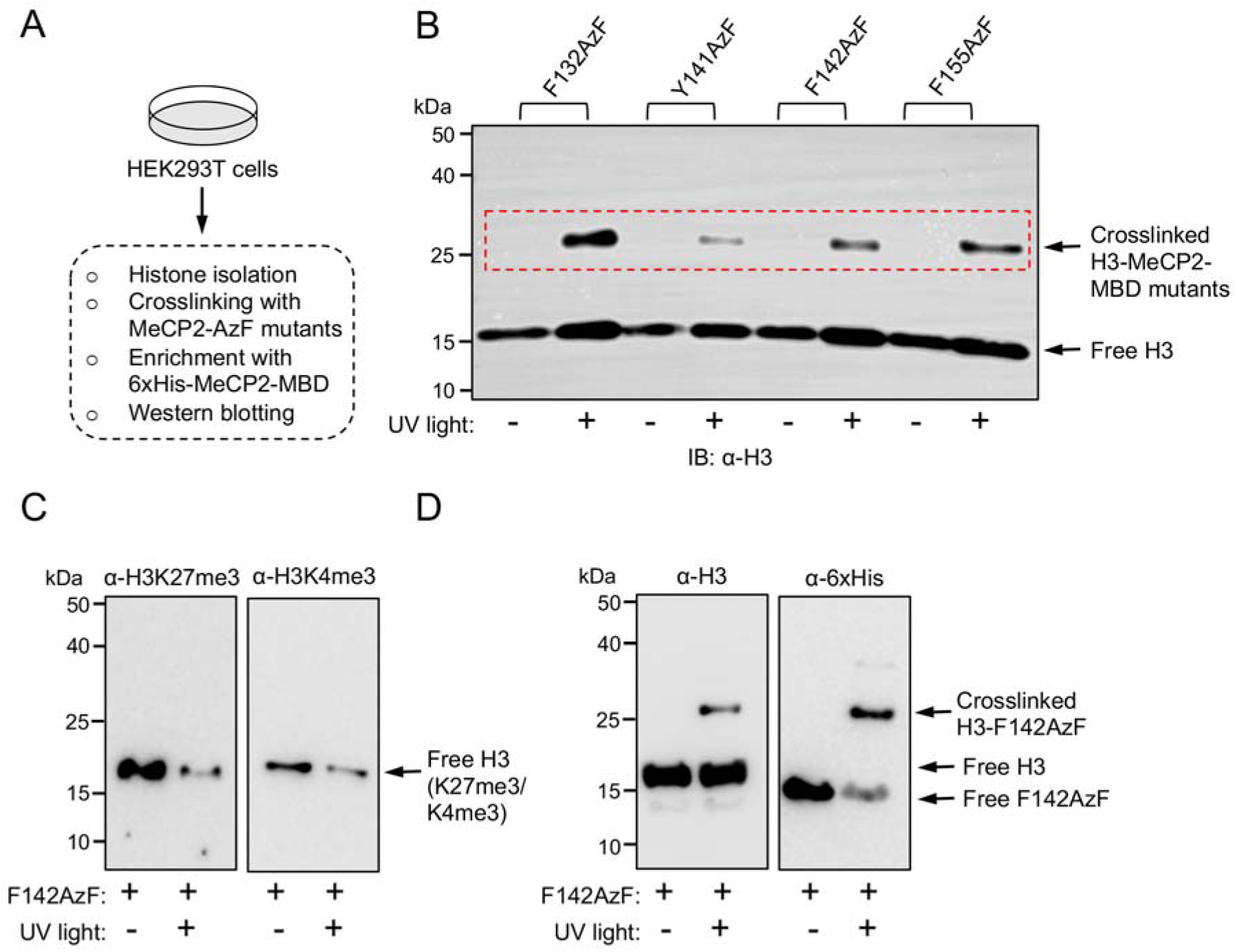
Photocrosslinking of engineered MeCP2-MBD with histones. (A) Schematic representation of histone isolation and photocrosslinking with engineered MeCP2-MBD. (B) Western blot showing photocrosslinking of engineered MeCP2-MBD variants with histone H3. (C) Western blot showing specific crosslinking of the F142AzF mutant with methylated histone ligands H3K27me3 and H3K4me3. (D) Confirmation of F142AzF crosslinking with histone H3 by immunoblotting using anti-H3 and anti-6xHis antibody.

For subsequent experiments, we focused on the F142AzF variant, since our previous study showed that, among the five aromatic cage mutants of MeCP2-MBD, only F142A retains partial histone-binding affinity (30). To determine whether MeCP2-MBD-F142AzF could crosslink with tri-methylated histones, we probed the crosslinked products with H3K4me3 and H3K27me3 antibodies, which recognize its two known ligands. Immunoblotting revealed a reduction in signal intensity for H3K4me3 and H3K27me3 in the UV-treated samples, with no detectable crosslinked bands (Figure 2C). However, when the same samples were re-probed with anti-H3 and anti-6xHis antibodies, high-molecular-weight bands corresponding to crosslinked complexes were observed (Figure 2C). These results demonstrate that MeCP2-MBD-F142AzF efficiently crosslinks with tri-methylated histone ligands, and the lack of detectable H3K4me3 and H3K27me3 signals likely reflects epitope masking within the crosslinked complex, consistent with these modifications occupying the ligand-binding pocket of the engineered reader.

### Photocrosslinking of engineered MeCP2-MBD with the cellular proteome

The successful crosslinking of F142AzF with its methylated histone ligands prompted us to investigate proteome-wide interacting partners of MeCP2. HEK293T cell lysates were incubated with MeCP2-MBD-F142AzF and irradiated with 365 nm UV light for 15 minutes. Crosslinked complexes were enriched using Ni-NTA beads and extensively washed to remove non-specific binders. The eluted proteins were resolved by SDS–PAGE and analyzed by western blotting with an anti-His antibody. Multiple high-molecular-weight bands appeared specifically in the UV-treated samples (Figure 3A), indicating successful crosslinking of the engineered MeCP2-MBD to putative interacting partners in the cellular lysate. No significant enrichment was observed in samples not exposed to UV light.

**Figure 3.**
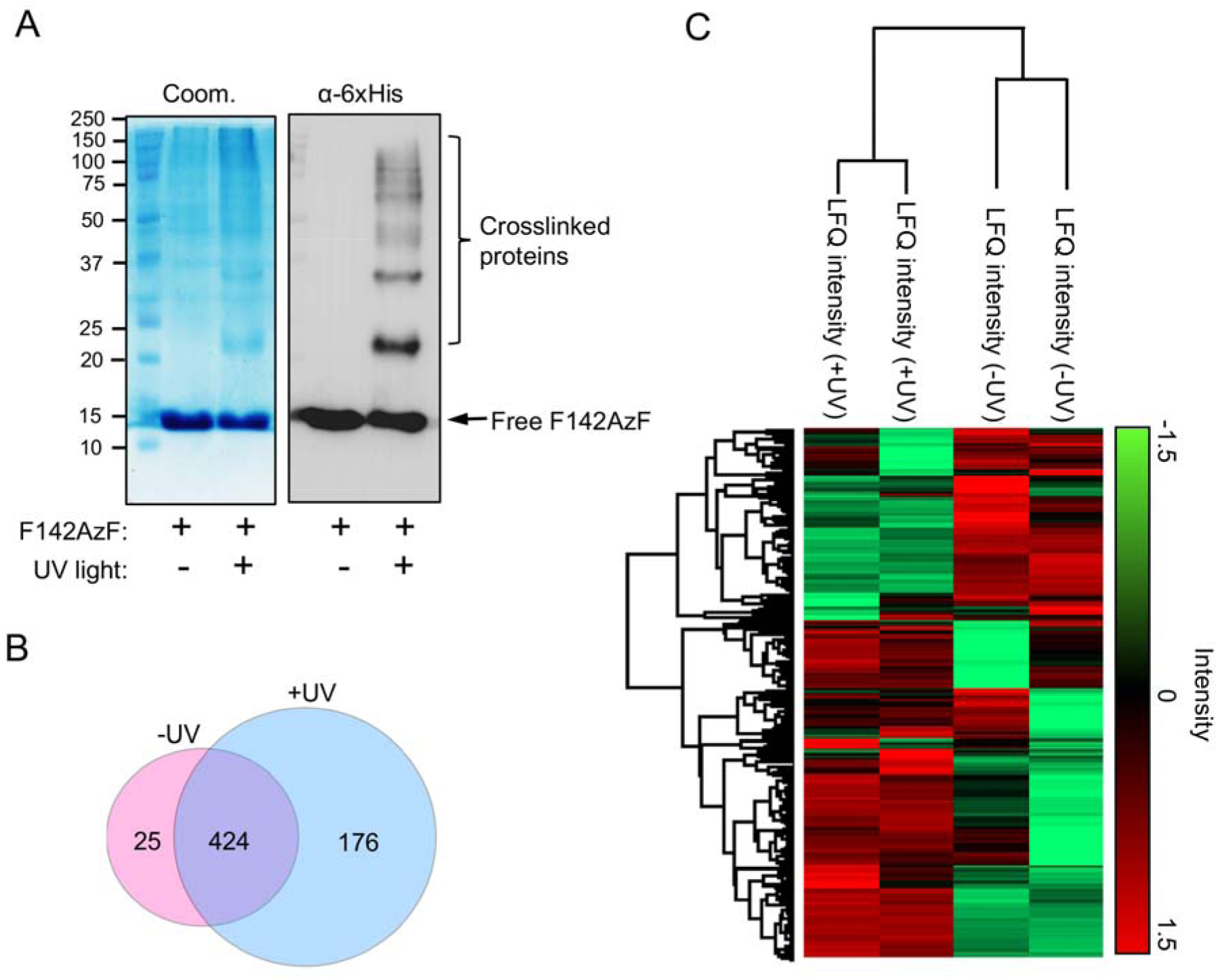
Photocrosslinking of F142AzF with cellular proteome. (A) Coomassie-stained gel and Western blot showing the enriched MeCP2 interactome pulled down from HEK293T cell extracts with or without UV irradiation. (B) *Venn diagram* showing number of proteins identified exclusively in −UV and +UV treated samples as well as those common to both conditions. (C) Heat map representing LFQ intensities of the identified proteins across both biological replicates for each condition.

To identify these interacting partners, we performed mass-spectrometry-based proteomic analysis. Enriched proteins were subjected to in-solution trypsin digestion followed by characterization using liquid chromatography-tandem mass spectrometry (LC-MS/MS). Two independent biological replicates of both UV-treated and untreated samples were analyzed to ensure reproducibility and confidence in the dataset. Label-free quantification (LFQ) was carried out using the MaxQuant software suite, and the resulting “proteinGroups.txt” file was further processed in Perseus. Proteins annotated as potential contaminants or reverse hits were removed. In total, 625 proteins were identified; among these, 25 were detected exclusively in the untreated samples, 176 were unique to the UV-treated condition, and 424 were common to both (Figure 3B). To assess reproducibility between replicates, a heatmap based on LFQ intensities was generated, demonstrating a high degree of consistency across samples (Figure 3C).

### Identification of significantly enriched MeCP2 interactome

To identify proteins significantly enriched upon photocrosslinking of the engineered MeCP2-MBD, LFQ intensities from UV-treated and untreated samples were compared in Perseus. A volcano plot was generated using a two-sample *t*-test with an FDR cutoff of 0.05 and an S_0_ value of 0.1 (Figure 4A and Table S1). Several proteins were significantly enriched in the UV-treated condition, indicative of specific MeCP2-mediated crosslinking events. Proteins meeting the criteria of *p* ≤ 0.05 and ∼2-fold enrichment were classified as high-confidence MeCP2 interactors (Figure 4A and Table S1).

**Figure 4.**
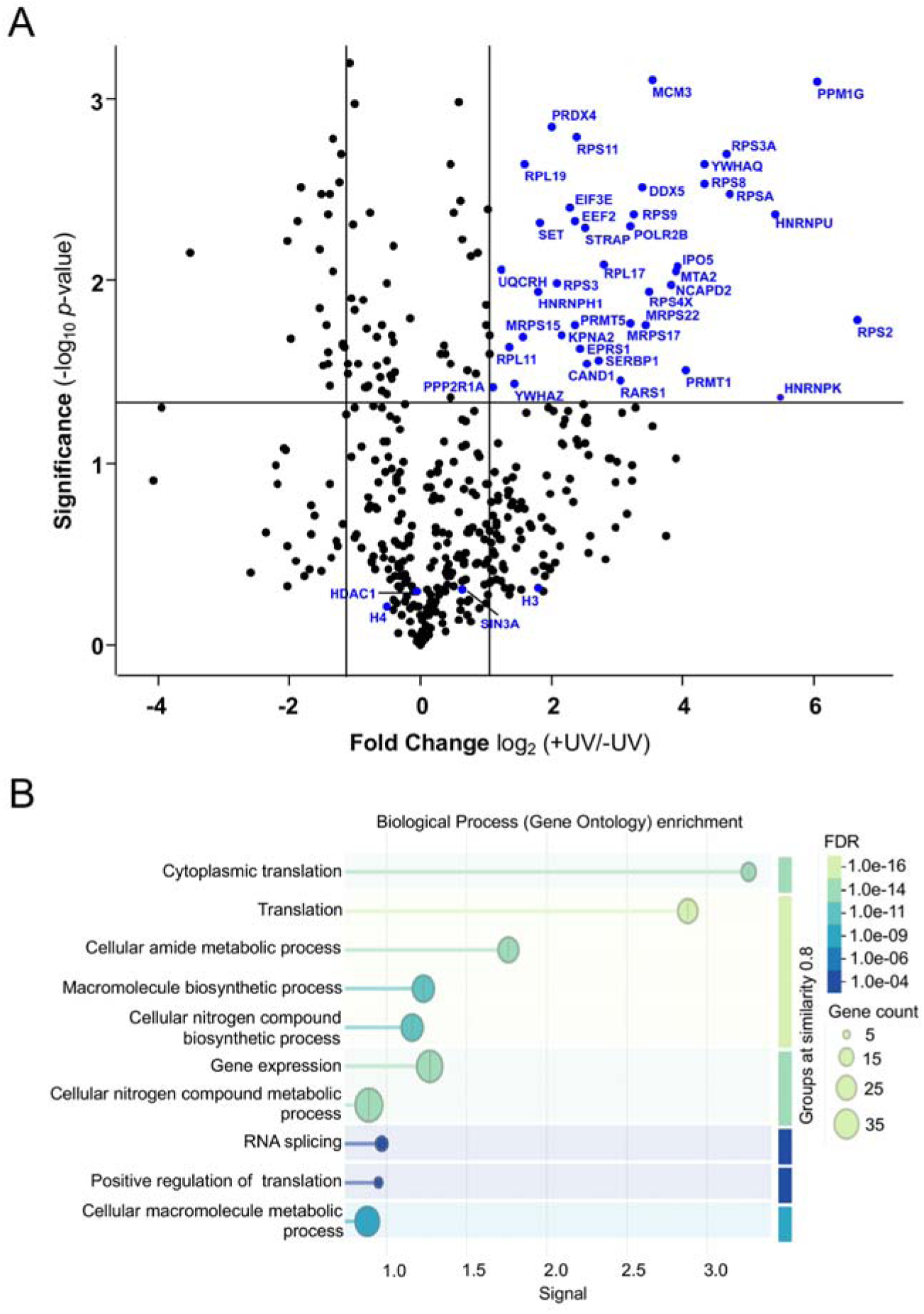
Identification and functional characterization of high-confidence MeCP2 interactome. (A) A *volcano plot* represents potential interacting partners of MeCP2-MBD identified by proteomic analysis. (B) Gene Ontology (GO) enrichment analysis of the significant MeCP2-interacting proteins.

To gain insight into the functional categories represented by these interacting partners, the significantly enriched proteins were subjected to Gene Ontology (GO) enrichment analysis (Figure 4B). The enriched GO terms were predominantly associated with chromatin and transcriptional regulation (POLR2B, SET, MTA2, PRMT1, PRMT5, PPM1G), translation regulation (EIF3E, EEF2, EPRS1, RARS1), protein transport (IPO5, KPNA2), RNA processing and splicing (HNRNPU, HNRNPK, HNRNPH1, DDX5, SERBP1, STRAP), mitochondrial translation (MRPS15, MRPS17, MRPS22), and ribosomal proteins (RPS/RPL families). Notably, several biologically important interacting partners were uniquely detected under the +UV condition, including TP53 (a key transcription factor and tumor suppressor), SIRT1 (an NAD⁺-dependent deacetylase regulating transcription and stress responses), WDR26 (a scaffolding protein involved in signal transduction), G3BP1 (an RNA-binding protein and core stress granule component), and PRKACA (the catalytic subunit of protein kinase A). Together, these findings indicate that MeCP2 engages a diverse network of chromatin-associated, transcriptional, and post-transcriptional regulators within the cellular environment, underscoring its multifaceted regulatory roles.

## Discussion

The MBD of MeCP2 is known to recognize several tri-methylated histone marks, including H3K4me3, H3K9me3, H3K27me3, and H3K36me3 (30). Among these, the interaction between MeCP2-MBD and H3K27me3 has been shown to play a key role in gene expression regulation (33). In our previous work, we defined the tri-methyllysine–binding pocket within MeCP2-MBD, formed by an aromatic cage that mediates recognition of methylated histone ligands (30). Building on this foundation, we developed a photocrosslinking-based chemoproteomic strategy to map the aromatic cage–dependent interactome of MeCP2-MBD. By site-specifically incorporating the photocrosslinkable amino acid AzF into the methyllysine-binding aromatic cage, we captured both histone and non-histone binding partners of MeCP2.

In this study, we showed that phenylalanine and tyrosine residues within the aromatic cage can be substituted with AzF without disrupting MeCP2’s ability to recognize tri-methylated histones. All engineered MeCP2-MBD-AzF variants exhibited robust crosslinking with histone H3. Among these, MeCP2-MBD-F142AzF was chosen for cellular proteome crosslinking, based on prior mutational analyses demonstrating that F142A partially retains binding affinity for methylated histones (30). Proteomic profiling of MeCP2-MBD-F142AzF crosslinked complexes from HEK293T cell lysates identified numerous high-confidence interacting partners, demonstrating that aromatic cage engineering enables efficient capture of dynamic, methyllysine-dependent interactions.

Gene Ontology analysis revealed that high-confidence MeCP2 interactors participate in chromatin organization, transcriptional regulation, RNA processing, translation, and metabolic pathways. Chromatin-associated proteins such as MTA2 (a subunit of the NuRD complex), PRMT1/5 (arginine methyltransferases), and SET (a lysine methyltransferase) suggest that MeCP2 cooperates with multiple chromatin-modifying enzymes to regulate gene activity. Beyond chromatin regulation, we also observed extensive interactions with RNA-binding and processing factors including HNRNPU, HNRNPK, DDX5, and SERBP1 indicating a broader role for MeCP2 in post-transcriptional regulation. The enrichment of translation-related proteins (EIF3E, EEF2, EPRS1) and mitochondrial ribosomal components (MRPS15, MRPS17, MRPS22) further supports the idea that MeCP2 may function as an integrative node linking transcriptional, post-transcriptional, and metabolic processes.

In addition to identifying new interactors, our photocrosslinking strategy also recovered several previously reported MeCP2-associated proteins, underscoring the robustness of the approach. Histones H3 and H4, which have long been established as MeCP2 binding partners (33–36), were readily detected in our dataset. Likewise, the corepressors SIN3A and HDAC1 components of the canonical MeCP2 repressive complex (10,11) were also identified. Recent work has shown that MeCP2 directly interacts with RNA polymerase II at the transcription start sites of active genes (37); consistent with this, POLR2B (RNA polymerase II subunit B) emerged as a high-confidence interactor in our analysis. Collectively, these results demonstrate that our photocrosslinking-based chemoproteomic strategy effectively captures both established and previously unrecognized MeCP2-associated proteins. This approach provides a powerful and reliable platform for mapping MeCP2’s interactome with high confidence, offering new insights into the diverse biological pathways regulated by this essential epigenetic reader.

## Conclusion

In summary, our study demonstrates that the aromatic cage of MeCP2-MBD is not only essential for recognizing methylated histone marks but also plays a key role in mediating interactions with a broad spectrum of non-histone proteins. The diversity of interaction partners uncovered here supports the view that MeCP2 functions as a multifunctional chromatin-associated regulator that coordinates multiple layers of gene expression from chromatin remodeling and transcriptional control to RNA processing and translation. The wide range of enriched proteins also suggests that MeCP2 may act as an integrative hub that couples epigenetic information with cellular signaling and metabolic pathways. Importantly, the chemical biology strategy developed in this work combining aromatic cage engineering with photocrosslinking-based proteomic profiling establishes a generalizable platform for mapping methyllysine-dependent interactions of other chromatin reader proteins. Future studies will be essential to elucidate the functional consequences of these interactions in biologically relevant contexts, including neuronal differentiation and tumorigenesis.

## Materials and methods

### General materials, methods, and equipment

All of the plasmids for bacterial expression were obtained as gifts from individual laboratories or purchased from addgene. Mutagenic primers were obtained from Sigma-Aldrich (Table S2). Commercially available competent bacterial cells were used for protein expression and mutagenesis. HEK293T cells were obtained from American Type Culture Collection (ATCC) and used following the manufacturer’s protocol. All of the antibodies used in this study were purchased from established vendors and used following the manufacturer’s protocol. 4-azido-L-phenylalanine (AzF) was a kind gift from K. Islam (University of Pittsburgh, Pittsburgh, PA) or purchased from SciTech and characterized as reported previously (38).

### Protein-peptide docking

Protein–peptide docking was performed using the HADDOCK2.4 web server (39) using method as previously described (30). The 3D structure of the MeCP2-MBD domain was obtained from the PDB (PDB code 6OGK), and the peptide representing H3(21–36)K27me3 was built in an extended conformation using PyMOL. Both the protein and peptide structures were uploaded to the server, and the active site residues within the protein’s putative binding site were specified. The docking process was carried out in three stages: 1) rigid-body energy minimization (it0), 2) semi-flexible simulated annealing refinement (it1), and 3) final refinement in explicit solvent (water refinement). The resulting top clusters were ranked based on the HADDOCK score, a weighted sum of van der Waals interactions, electrostatics, desolvation energy, and restraint violations. The best-ranked cluster, characterized by the lowest average HADDOCK score, favorable interface RMSD, and a high buried surface area, was selected for further analysis. All models were visualized and assessed using PyMOL.

### Mutagenesis, expression, and purification of the MeCP2-MBD domain

The C-terminal 6xHis-tagged MeCP2-MBD domain bacterial expression construct pNIC-CH was obtained from addgene (a gift from Cheryl Arrowsmith, addgene catalog no. 162259). Protein expression and purification were performed using previously described methods (30). The wild-type MeCP2-MBD plasmids was transformed into One Shot BL21 star (DE3) *E. coli* competent cells (Invitrogen, catalog no. C601003) using pNIC-CH kanamycin-resistant vector. A single colony was picked up and grown overnight at 37 °C in 10 mL of Luria-Bertani (LB) broth in the presence of 50 μg mL^−1^ kanamycin. The culture was diluted 100-fold and allowed to grow at 37 °C to an optical density (OD_600_) of 1.0. Protein expression was induced overnight at 17 °C with 0.5 mM isopropyl β-D-1-thiogalactopyranoside in an Innova 44 Incubator shaker (New Brunswick Scientific). Proteins were purified as follows. Harvested cells were resuspended in 15 mL of lysis buffer [50 mM HEPES (pH 7.5), 500 mM NaCl, 5 mM β-mercaptoethanol, 5% glycerol, 25mM imidazole, lysozyme, DNase, and 1:200 (v/v) Protease Inhibitor Cocktail III (Calbiochem)]. The cells were lysed by pulse sonication and centrifuged at 13000 rpm for 30 min at 4°C. According to the manufacturer’s instructions, the soluble extracts were subjected to Ni-NTA agarose resin (QIAGEN, catalog no. 30210). After 20 volumes of wash buffer [50 mM HEPES (pH 7.5), 300 mM NaCl, 5 mM β-mercaptoethanol, 5% glycerol, and 25 mM imidazole] had been passed through the column, proteins were eluted with a buffer containing 50 mM HEPES (pH 7.5), 300 mM NaCl, 5 mM β-mercaptoethanol, 5% glycerol, and 250 mM imidazole. Proteins were further purified by gel filtration chromatography (Superdex-75) using an AKTA pure FPLC system (GE Healthcare) in a buffer containing, 50 mM HEPES (pH 7.5), 200 mM NaCl, and 5% glycerol. Purified proteins were concentrated using an Amicon Ultra-10k centrifugal filter device (Merck Millipore Ltd.), and the concentration was determined using a Bradford assay kit (Bio-Rad Laboratories) with bovine serum albumin as a standard. The proteins were aliquoted and stored at −80 °C until use.

Site-directed mutagenesis was performed to generate the MeCP2-MBD amber codon (TAG) mutants F132TAG, Y141TAG, F142TAG and F155TAG using the QuikChange Lightning Site-Directed Mutagenesis Kit, and the resulting mutant plasmids were confirmed by DNA sequencing. To express MeCP2-MBD variants carrying 4-azido-L-phenylalanine (AzF) at site-specific positions, MeCP2-MBD amber variants (F132TAG, Y141TAG, F142TAG and F155TAG) and pEVOL-pAzF (a kind gift from K. Islam, University of Pittsburgh, addgene catalog no. 31186) were co-transformed into One Shot BL21 Star (DE3) *E. coli* competent cells. After transformation, cells were recovered in 200 μL of SOC medium and incubated in a 37 °C shaker for 1 h at 225 rpm before being plated on a dual antibiotic LB Miller agar plate containing 50 μg mL^−1^ kanamycin and 35 μg mL^−1^ chloramphenicol. Protein expression and purification were performed using previously described methods (40). Briefly, a single colony was picked up and inoculated into 5 mL of LB Miller broth in the presence of 50 μg mL^−1^ kanamycin and 35 μg mL^−1^ chloramphenicol and grown overnight in a 37 °C shaker at 225 rpm. This overnight culture was centrifuged (Eppendorf Centrifuge 5910 R, S-4xUniversal swing-bucket rotor) at 1000g (2158 rpm) for 10 min at room temperature, and the supernatant (LB Miller broth) was discarded. The collected cell pellet was then resuspended in 1 mL of M9 medium and used to inoculate 500 mL of GMML medium supplemented with 50 μg mL^−1^ kanamycin and 35 μg mL^−1^ chloramphenicol. Cells were allowed to grow at 37 °C in an incubator shaker (225 rpm) to an optical density (OD_600_) of 1.0. The unnatural amino acid (AzF) was prepared by dilution in 5 mL of sterilized deionized water, and that mixture was added to the bacterial culture to a final concentration of 1 mM. Then the cells were cooled to 17 °C and allowed to grow for 30 min; at this stage, AzF-specific aminoacyl-tRNA synthetase expression was induced with 0.05% (w/v) arabinose and allowed to shake for an additional 30 min at 17 °C. Finally, MeCP2-MBD protein expression was induced by the addition of 0.5 mM IPTG and cells were allowed to grow for 20 h at 17 °C. The MeCP2-MBD mutants were purified as described above while minimizing the exposure to ambient light.

### Mammalian cell culture

HEK293T cells were cultured in DMEM supplemented with 10% fetal bovine serum and a penicillin/streptomycin antibiotic cocktail in a humidified atmosphere containing 5% CO_2_. After 20 hours, the medium was removed, and the cells were rinsed with cold phosphate-buffered saline (PBS). The harvested cells were washed again with cold PBS and then frozen as a dry pellet at −80 °C.

### Acid extraction of histones

Histones were extracted from HEK293T cells using the acid-extraction protocol as previously described (41). Briefly, frozen cell pellets from T75 flask of HeLa cells were resuspended in 1 ml of hypotonic lysis buffer (10 mM Tris-HCl pH 8.0, 1 mM KCl, 1.5 mM MgCl_2_, 1 mM DTT, 1 mM PMSF, and protease inhibitor cocktail) and incubated for 30 min on rotator at 4 °C. Samples were then centrifuged (10,000 x g, 10 min at 4 °C), and the supernatant was discarded entirely. The nuclei resuspended in 800 μL of 0.4 N H_2_SO_4_, vortexed intermittently for 5 min, and further incubated at 4 °C on a nutator for overnight. The nuclear debris was pelleted by centrifugation (16000 x g, 10 min at 4 °C), and the supernatant containing histones were collected. The histones were precipitated by adding 264 μL TCA (Trichloroacetic acid) drop by drop to histone solution and invert the tube several times to mix the solutions and incubated the samples on ice for 30 min. Finally, histones were pelleted by centrifugation (16000 x g, 10 min at 4 °C), and the supernatant was discarded. The histones pellet was washed twice with ice-cold acetone, followed by centrifugation (16000 x g, 5 min at 4 °C), and carefully removed the supernatant. The histone pellet was air-dried for 20 min at room temperature and subsequently dissolved in an appropriate volume of ddH2O and transferred into a fresh tube. The aliquoted histones were stored at -80 °C until further use.

### Photocrosslinking with histones and western blotting

Photocrosslinking with histone extracts was performed by methods as previously described with slight modifications (38,40). Briefly, 20-25 μg histones (extracted from HEK293T cells) were incubated with 50 μM MeCP2 MBD-AzF mutants (F132AzF, Y141AzF, F142AzF and F155AzF) one at a time in 100 μL binding buffer (50 mM HEPES pH 7.5, 150 mM NaCl, 0.001% Tween 20) for 30 min at 4°C on a rotator. After incubation, the samples were split into two 50 μL volumes in clear PCR tubes (Axygen). The sample-containing PCR tubes were placed into semimicro visible cuvettes (300−900 nm, Eppendorf, catalog no. 0030079353) filled with ice cubes to maintain the temperature and irradiated with ultraviolet light (365 nm, 8 W lamps, model ECX-F20.L V1, Vilber Lourmat) for 3 x 10 min at 4 °C. Negative samples were not subjected to ultraviolet (UV) irradiation and stored in the dark at 4 °C. After irradiation, the samples were placed in 1.5 mL tubes having 50-60 μL of Ni-NTA agarose beads (QIAGEN) pre-equilibrated with the buffer [50 mM HEPES (pH 7.5), 150 mM NaCl, and 0.001% Tween 20], and the samples were incubated for 1 h at 4 °C on a rotator. The uncrosslinked histones were removed by washing the samples with washing buffer [50 mM HEPES (pH 7.5), 500 mM NaCl, and 0.5% Triton X-100] five times at room temperature. The proteins were eluted with 30 μL of elution buffer containing 50 mM HEPES (pH 7.5) and 500 mM imidazole and incubated for 5 min at room temperature with intermittent mixing. The eluted proteins were mixed with 10 μL of 1x TruPAGE LDS sample buffer (Sigma, catalog no. PCG3009) and heat-denatured at 95 °C for 10 min. Finally, the cross-linked proteins were separated on a 12%-SDS-PAGE gel and transferred onto a 0.45 μm PVDF membrane at a constant voltage of 80V for 1 hr. at 4°C. The membrane was rinsed in TBST buffer (50 mM Tris pH 7.4, 150 mM NaCl, and 0.1% Tween-20) and blocked for an hour at room temperature (RT) in 5% milk buffer prepared in TBST. Immunoblotting was performed with the following primary antibodies: anti-H3 (Cell Signaling Technology, catalog no. 9715), anti-H3K27me3 (Cell Signaling Technology, catalog no. 9733), anti-H3K4me3 (Invitrogen, catalog no. 703954) and anti-6xHis (Invitrogen, catalog no. MA121315) overnight at 4°C. The membranes were washed with TBST buffer thrice at RT for five minutes each. The blots were then incubated with the HRP conjugated secondary antibodies Goat anti-Rabbit IgG (Invitrogen, catalog no. 31466) or Goat anti-mouse IgG (Invitrogen, catalog no. 31431) with 5% nonfat dry milk, dilution 1:5000 in TBST. The membranes were rewashed with TBST buffer thrice at RT for five minutes each. Protein bands were visualized by chemiluminescence using SuperSignal West Pico PLUS substrate (Invitrogen, catalog no. 34577) following the manufacturer’s protocol.

### Preparation of HEK293T cell lysate

The harvested HEK23T cells were resuspended in ice cold lysis buffer containing 25 mM Tris-HCl pH 7.4, 150 mM NaCl, 1% NP-40, 1 mM EDTA, 5 % glycerol (Pierse IP lysis buffer, catalog no. 87787), and supplemented with 1X Pierse protease inhibitor cocktail and incubated on ice for 5 min with periodic mixing. Cell lysate was centrifuged at 13, 000 x g for 10 min at 4 °C to pellet the cell debris. The supernatant was transferred into fresh microcentrifuge tube and the protein concentration was determined by Bradford assay (Bio-Rad Laboratories). The supernatant containing cell lysate was used for photocrosslinking, western blotting, and subsequent proteomic studies.

### Photocrosslinking with HEK293T cell lysate and western blotting

For photo-cross-linking studies, ∼1.0 mg of HEK293T cell lysate was incubated with 50 μM MeCP2-MBD-F142AzF mutant in a buffer containing 50 mM HEPES pH 7.5, 150 mM NaCl, 0.001% Tween 20. After 1 h of incubation at room temperature, the samples were subjected to UV irradiation at 365 nm for 30 min at 4 °C. Negative controls were not subjected to UV exposure. Samples were then bound to Ni-NTA agarose resin and incubated for 1h at 4 °C with gentle rotation. To remove un-crosslinked proteins, present in cell lysates, samples were washed 5 times with washing buffer (50 mM HEPES pH 7.5, 400 mM KCl, 5% Triton X-100). The cross-linked proteins were eluted in 30 μL elution buffer containing 50 mM HEPES pH 7.5, 500 mM imidazole and incubated for 5 min at room temperature with intermittent mixing. The eluted proteins were separated on a 4-15% Mini-PROTEAN TGX precast SDS-PAGE gel (Bio-Rad, #4561085) and transferred onto a 0.45 μm PVDF membrane at a constant voltage of 80V for 1.5 hr. at 4°C. The membrane was rinsed in TBST buffer (50 mM Tris pH 7.4, 150 mM NaCl, and 0.1% Tween-20) and blocked for an hour at room temperature (RT) in 5% milk buffer prepared in TBST. Immunoblotting was done with anti-6xHis tag antibody (Invitrogen, catalog no. MA121315) overnight at 4°C. The membrane was washed with TBST buffer thrice at RT for five minutes each. The blots were then incubated with the HRP conjugated secondary antibody Goat anti-mouse IgG (Invitrogen, catalog no. 31431) with 5% nonfat dry milk, dilution 1:5000 in TBST. The membrane was rewashed with TBST buffer thrice at RT for five minutes each. Protein bands were visualized by chemiluminescence using SuperSignal West Pico PLUS substrate (Invitrogen, catalog no. 34577) following the manufacturer’s protocol.

### LC-MS/MS analysis

In-solution trypsin digestion was carried out by following the standard protocol. Proteolytic peptides from in-solution trypsin digestion were analyzed by a nanoflow reverse-phased liquid chromatography tandem mass spectrometry (LC-MS/MS). Tryptic peptides were loaded onto a PepMap RSLC C18 column (2μm, 100Å x 50 cm) of the LC system (Q-Exactive Plus Biopharma, Thermo Scientific). Chromatographic separation was performed using a binary solvent system (solvent A: water with 0.1% formic acid; solvent B: acetonitrile and water in 85:15 ratios with 0.1% formic acid). MaxQuant v2.6.8.0 software was used for protein identification and quantitation. Mass spectrometry data were searched against UniProt-proteome-UP000005640 using Andromeda. Protein identifications were reported with a false discovery rate (FDR) of 1%. We used MaxQuant to calculate iBAQ, a measure of protein abundance.

### Data processing and statistical analysis in MaxQuant-Perseus

Perseus v2.1.4.0 software was used to further process the proteomics data. The ‘proteinGroups.txt’ file generated by MaxQuant was used as input. Proteins identified ‘only by sites’ as well as those annotated as ‘reverse’ or ‘potential contaminants’ were removed. Biological replicates were grouped into single experimental conditions and LFQ intensities were log_2_ transformed. Rows containing missing values were filtered with a 75% valid-value threshold. Missing values were imputed from a normal distribution to simulate signals of low-abundance proteins. Protein annotations were added for *Homo sapiens* from the integrated Perseus annotation files. Heat map was generated using z-score normalized LFQ intensities to visualize clustering between biological replicates. Volcano plot was constructed to identify differentially abundant proteins, using a with FDR cutoff of 0.05 and an S_0_ parameter of 0.1.

## Acknowledgments

The authors thank infrastructural facilities supported by IISER Kolkata (Ministry of Education), Government of India.

## Author Contributions

B.S. conceived the ideas. J.P., and B.S. designed the experiments. J.P. performed the experiments. J.P., and B.S. analyzed data. J.P., and B.S. prepared original draft. B. S. writing-reviewing and editing. B.S. supervision, B.S. resources. B.S. project administration. B.S. funding acquisition.

## Funding and additional information

This work was supported by grants from ANRF (CRG/2022/005242), SERB (EEQ/2020/000149), DBT Ramalingaswami Fellowship (BT/RLF/Re-entry/56/2018), and an intramural grant from IISER Kolkata (Ministry of Education), Government of India, to B.S. The authors also acknowledge the Prime Minister’s Research Fellowship (PMRF) program for providing doctoral fellowship and research grant to Jyotirmayee Padhan (PMRF ID-0501974), Government of India.

## Abbreviations

MeCP2: Methyl-CpG binding protein 2
MBD: methyl-CpG-binding domain
AzF: 4-azido-L-phenylalanine
LFQ: Label-free quantification (LFQ)

